# Rational Design of Peptide Vaccines for the Highly Lethal Nipah and Hendra Viruses

**DOI:** 10.1101/425819

**Authors:** Sumanta Dey, Proyasha Roy, Tathagata Dutta, Ashesh Nandy, Subhash C Basak

## Abstract

The Nipah virus disease is a lethal infection that has led to 40% to 75% fatalities in Malaysia, Bangladesh and India. The reports of human-to-human transmission documented in Bangladesh has raised the specter of pandemic potential and has caused the World Health Organization to list the Nipah virus as one of the pathogens to be considered for development of drugs and vaccines on urgent basis, neither of which exist against the Nipah virus as of now, although many proposals have been made and trials initiated. Given that there are established country-specific differences in the virus’ effects and fatalities, meeting the sudden need for a vaccine in case of an epidemic will require design, development and preparation for a peptide vaccine. Thus, we propose a protocol for creating peptide vaccines that can be tailor-made for these specific countries, an approach which is being advocated for the first time. Here, we analyze the surface proteins, Fusion protein and Glycoprotein, of the strains currently affecting the three countries on a large scale and determine the specific country-based epitope differences.

## Introduction

A lethal zoonotic paramyxovirus with a possibility of pandemic potential is affecting wide swathe of tropical populations from Philippines to India to Madagascar [1]. First isolated from a patient in Kampung Sungai Nipah village of Malaysia in 1998 [2], the Nipah virus (NiV) caused fatalities in Malaysia among people in the pig trades. Fruit bats, also known as flying foxes, of the *Pteropus* genus have been identified as the natural reservoirs of the virus, with pigs, horses and other domesticated animals as carriers of the disease. Culling of over a million pigs halted the progress of the infections in Malaysia, but the disease next struck in Bangladesh in 2001 where the infection is reported to have spread through contaminated fruits and date palm sap [3]. The infection has become since an annual occurrence there with occasional infections in neighbouring India [4], and lately in Kerala [5], far removed from Bangladesh. In 2014, the disease was also identified in the Philippines where a number of horses, and some people in contact with the horses, died from the infections [6]. Similar viruses, not yet human infecting, have been found in bats in Africa from Madagascar to Ghana [1], which indicates that the geographical range of the infectious disease could expand in future.

Nipah was determined to be closely related to the Hendra virus (HeV), first identified in horses in Australia, the two viruses forming a subspecies, the Henipavirus, in the Paramyxoviridae family, although the HeV is not known to infect humans the way the NiV has been seen to do, but has led to some deaths in Australia. The virus causes respiratory and encephalitic disorders in symptomatic individuals leading to death in severe cases; in Malaysia the case-fatality ratio (CFR) was about 40%, but in Bangladesh and India the CFR was between 70% and 100%. The pathogenicity of the two strains of NiV seem to be different, with the Bangladesh strain being more lethal. This was also borne out by experiments with ferrets [7] and African green monkeys [8], showing that any post-infection therapies need to be specific to the strains involved.

The disease is transmitted through physical contact with bodily secretions and excretions of infected animals, but more importantly, human-to-human infections through exhaled droplets from respiratory organs have been known to occur in about 50% of the cases in the Bangladesh infections [9]. So far with NiV infections being in localized remote villages, the current spread of the highly lethal viral infections is as yet limited in numbers. However, RNA viruses being prone to rapid mutational changes, it could be a matter of time before the virus may mutate to a more efficient human-to-human transmissibility, which, coupled with the density of population in these regions and with quite fluid human mobility, could lead the infection to reach pandemic proportions. This possibility has caused the CDC to classify NiV and HeV as Bio-Safety Level 4 viruses [10]. The NIAID in its classification of biodefense research has categorized the Nipah virus as a Category C emerging pathogen with potential for high morbidity and mortality rates, and major health impact [11]. Nipah is also on the World Health Organization research and development list of viruses that require urgent attention alongside Ebola, Zika, MERS, Lassa and Crimean-Congo hemorrhagic fever.

The recent incidences of Nipah virus infections in Kerala thus necessitates accelerated research in development of therapeutics and preventives against the Nipah and Hendra viruses. The Henipavirus is an enveloped, negative-sense RNA virus, approximately 18,250 nt long and consists of six genes: N (nucleoprotein), M (matrix), P (phosphoprotein), F (fusion), G (glycoprotein), and L (polymerase). The virus attaches itself to the Ephrin-B2 (EFNB2) receptor on host cell surface mediated by a phenylalanine side chain on EFNB2 that fits snugly into a hydrophobic pocket on the viral protein [12]. Currently, no drugs or vaccines exist for this virus, though many trials are in progress. Geisbert et al. [13] determined that an experimental human monoclonal antibody (mAb), m102.4, that targets the Ephrin-B2 and Ephrin-B3 receptor binding site of the HeV and NiV glycoprotein provides therapeutic remedy to African green monkeys when administered intravenously soon after infection before clinical symptoms develop. Dawes et al. [14] showed that a small molecule, Favipiravir (T-705), a purine analogue antiviral, administered subcutaneously in hamsters for 14 days fully protected the animals from a lethal dose of NiV and could therefore be considered as an antiviral treatment option. However, since these therapeutics are to be administered before clinical symptoms develop, their use in an epidemic scenario could present difficulties.

The major research emphasis has been on development of preventives through vaccination. As Satterfield et al. [15] has remarked, past experience with vaccine successes against other paramyxoviruses such as measles and mumps provide a reason to expect that a vaccine against the henipaviruses will also lead to positive results, although the extreme lethality of the Nipah virus remains a cause for concern. Of the various types of vaccines that can be developed, live attenuated viruses as vaccines are considered risky due to possibilities of reversion. Other approaches include live-vectored vaccine approaches, using vesicular stomatitis virus, Newcastle disease virus and others, and are under clinical trials. A subunit vaccine using soluble glycoprotein (sG) from the Hendra virus has been licensed to vaccinate horses in Australia against the HeV and can be tried against the Nipah virus since the two viruses have between 83% and 88% homology in the F and G proteins. Walpita et al. [16] reported a virus-like particle (VLP) vaccine composed of G, F and M proteins of the Nipah virus administered to hamsters provided complete protection to vaccinated animals against the viral challenge. However, there are very few licensed VLP vaccines; they are relatively more difficult to produce and cost considerations have kept other VLP vaccines such as ones against human papillomavirus out of reach of the majority of populations of developing countries.

An approach more suited to the economies and environments of the countries where the henipaviruses are prevalent is to consider peptide vaccines, although to date no such vaccines have been licensed for use on humans. Peptide vaccines are targeted against specific epitopes of the surface proteins of the virus and have many advantages [17,18]. In the current post-genomic era with improved tools of bioinformatics and growing knowledge of immune responses and immunogenetics, it is possible to contemplate vaccines tailored to suit each country, community, and perhaps individual (the personalized medicine approach), for most specific actions, in contrast with the “one size fits all” approach of traditional vaccines. Peptide vaccines can be designed *in silico* and the most promising candidates can be manufactured rapidly and tested, so such vaccines are can be relatively inexpensive compared to existing available methods of vaccine production and provide a ready means for combating emerging infectious diseases.

From this point of view, Hossain and Rubayet-Ul-Alam [19] designed B- and T-cell epitopes against the F protein of the Nipah virus and selected the epitopes with highest probability for the putative peptide vaccine against the Nipah virus. Saha et al. [20] considered the F, G and M proteins of the Nipah and Hendra viruses to determine a common B- and T-cell epitope-based vaccine that could elicit appropriate immune response against the henipaviruses. Bioinformatics tools were used to find the best epitopes and molecular docking simulations with MHC class-II (MHC II) and class-I (MHC I) molecules to confirm their antigenic potentiality and concluded that the epitopes from G and M, modified, proteins might constitute most effective target against the Nipah and Hendra viruses.

We had been advocating the design and development of peptide vaccines for some time now [21-25] and illustrated our protocol in Dey et al. (2017). The high virulence of the Nipah virus and its pandemic potential calls for a peptide vaccine strategy [26] for which we chose the surface situated G protein as the most appropriate target since it is responsible for attachment to the ephrin receptor in host cells. Since the Nipah viral sequence showed very strong, long stretches of conserved domains that matched HeV’s conserved stretches also, it was natural to club the two together to design a common vaccine that matched all our criteria. Using our protocol, we identified conserved segments on the viral protein which are sufficiently surface exposed, determined their epitope potential in the host population and tested for auto-immune threats. We report here on the results for the peptide targets for the Malaysian, Bangladeshi and Indian (Keralite) populations as a ready guide for developing appropriate vaccines. Such overall country/community-wise epitope search is, to the best of our knowledge, being reported for the first time.

## Materials and Methods

We downloaded all the available complete sequences of the Nipah virus glycoprotein (20) and the Hendra virus glycoprotein (15) and the Nipah glycoprotein structure, 2VWD, from the NCBI databases.

For numerical comparison of protein sequences, we used a sequence representation described earlier [27] where in a rectangular co-ordinate system of 20-dimensions each amino acid is associated with a particular direction and the protein sequence is plotted by taking successive steps in the directions dictated by the amino acids. This generates a plot in an abstract space, but we can define a weighted center of mass and a protein graph radius, *p_R_*. The special property of *p_R_* is that in each case when two sequences result in the same graph radius the sequences are found to be identical (28). Using a n-amino acid window (n < N, the total length of the protein sequence) and sliding it along a sequence we can compare the values of *p_R_* at each window position for a set of sequences; where the *p_R_* remain constant through all sequences for the same window position, or change very little, we can assume the peptide belonging to that window is reasonably well conserved. This property has been used in several instances previously [21-25, 27]. In this procedure specifically, the protein sequence is mapped using a window of 12 to 14 amino acids and sliding in increments of one amino acid (aa) at a time. At each stage the peptide graph radius, *p_R_*, is computed [27] and stored until the end of the sequence is reached. This is then repeated for each protein sequence of the selected group of sequences and then the entire lot is scanned at each mapped aa position to determine the number of different *p_R_* values amongst all the sequences. Since each *p_R_* value represents a particular peptide sequence, the number of these values at each aa position map out a protein variety profile along the sequence; a moving average over 12 windows’ values make for easy viewing on a plot whose minima indicate regions where variety, and by inference changes in amino acids, are relatively least indicating most conserved regions.

In this analysis we have used a peptide window size of 12 amino acids. In one of our earlier analysis with the neuraminidase protein [21] we had considered peptide lengths of 8, 10 and 12 amino acids; for rotavirus where mutational changes occur very frequently, we used window sizes of 12 to 14 amino acids [22]; 14 residue peptides were found to be more potent antigenically compared to smaller peptides [29] and we had used that size for analysis of influenza hemagglutinin [23]. We note that while peptides of 6 to 20 amino acids lengths are used in scanning for B-cell epitopes, peptides of 10+ residues may contain overlapping linear B-cell epitopes [30]; cytotoxic T-cell epitopes have a limited length (8—11 residues), helper T-cells use longer peptide lengths [31]. Considering these issues and the fact that the Nipah and Hendra viruses are RNA type which therefore is comparatively less stable, we fixed window length for this exercise at 12 amino acids.

The protein sequences are next mapped for solvent accessibility through an Average Solvent Accessibility (ASA) server such as SABLE or ITASSER or any other suitable web-based servers. Taking a moving average over 12 amino acids of solvent accessibility index, and averaging over several sequences for each virus type, this is mapped out to reveal by the crests on the graph the most solvent accessible regions.

Mapping both the graphs, peptide variability profile and solvent accessibility profile, together, we search for those regions where the protein variability is at a minimum and solvent accessibility is among the highest. These selected regions make up the first list of conserved solvent accessible segments of the protein.

To confirm that the regions identified in the previous step are indeed surface situated, we examine the identified regions in a protein 3D structure through PyMol software. Marking out the identified segments on one protein of the 3D structure will reveal how much of the segments lie on the surface; availability of nearby structures will reveal if these structures cover part or all of the selected segment(s).

Next, we consider which of the segments contains epitope regions both linear and discontinuous to elicit necessary immune response. For this we use the IEDB (Immune Epitope Database and Analysis Resource) server (http://tools.iedb.org/mhcii/) [32-34] as an epitope-prediction tool to predict MHC class I and MHC class II T-cell epitopes, and also for linear and discontinuous B-cell epitopes. In a large-scale evaluation of IEDB the prediction for MHC I epitopes have been seen to have accuracies of ~90%, while for MHC II this is less at 60 to 70% [35].

To see whether our selected peptide regions can act as epitopes and can able to elicit necessary immune response, we use another epitope-prediction tool ABCpred server (http://www.imtech.res.in/raghava/abcpred/ABC_submission.html) [36]. These tools indicate binding affinities of B-cell epitopes (with 66% accuracy).

We also needed to analyze the role of human MHC polymorphism in the Nipah disease severity. This is facilitated by downloading the available allelic combination and their frequency from Net allele Database link in IEDB for each country/community and compared the epitope potential of the viral protein.

Finally, we test the peptides to ensure they do not possess any autoimmune threats and are unique peptides by themselves. Each peptide segment is subjected to a protein-protein BLAST and only those that pass this test are retained. These segments then become our recommendations for peptide vaccine design subject to final verification in wet labs.

## Results and Discussions

Using a graphical representation and numerical characterization technique described in the Materials and Methods section, we computed *p_R_* values for the 35 Nipah and Hendra glycoprotein sequences using the sliding window technique for a window size of 12 amino acids. Fig. 1 shows the data smoothed out by a 12 aa moving average *p_R_* variability values. The figure also shows the ASA profile. From the graph the tentatively identified segments of amino acid stretches that show least variability with highest ASA values are marked by straight short green horizontal lines. The identified peptides are listed in Table 1.

**Figure 1.**
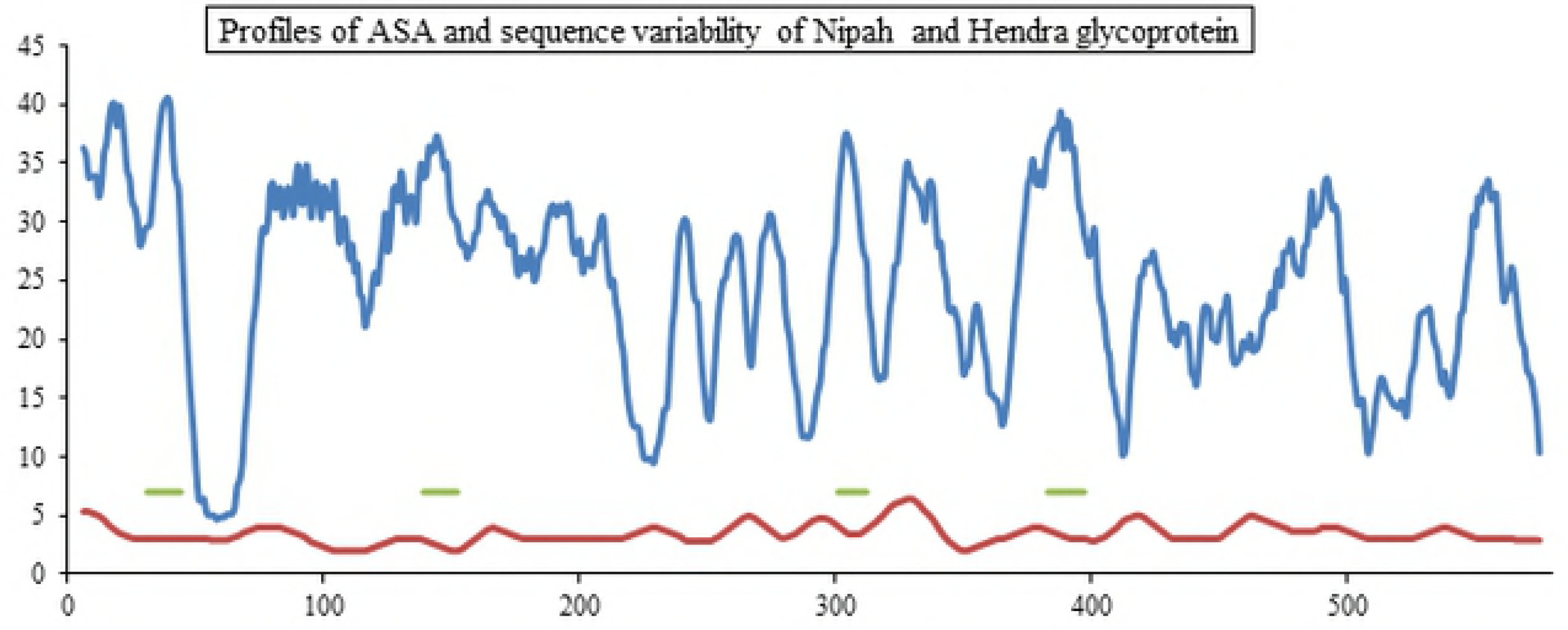
Profiles of average solvent accessibility (blue) in % and amino acid sequence variability (red) in numbers of the 35 Nipah and Hendra G glycoprotein sequences plotted against amino acid numbers. The short horizontal green lines are identified segments of the sequences where the amino acids are most conserved and have highest solvent accessibility.

**Table 1.**
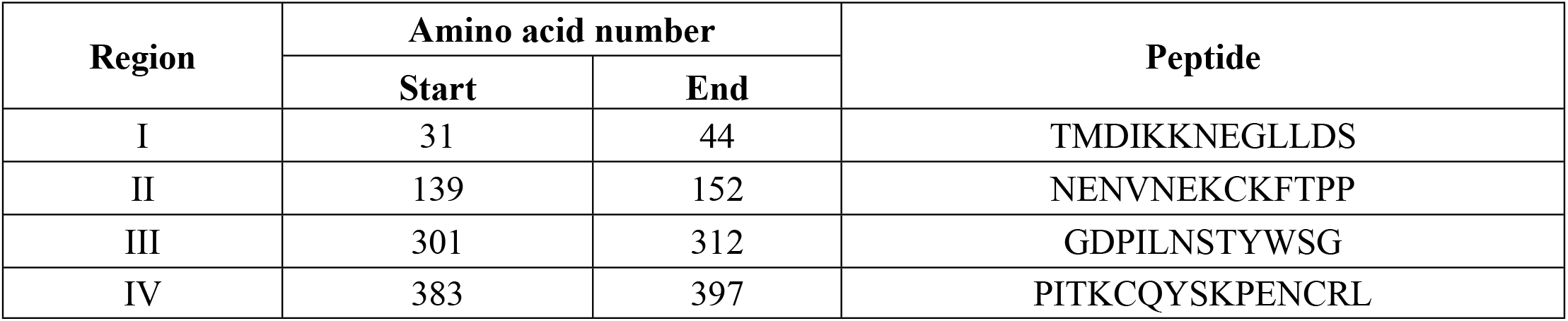
Regions with low amino acid variability and high solvent accessibility selected.

| Region | Amino acid number |  | Peptide |
| --- | --- | --- | --- |
|  | Start | End |  |
| I | 31 | 44 | TMDIKKNEGL LDS |
| II | 139 | 152 | NENVNEKCKFTPP |
| III | 301 | 312 | GDPI LNSTYWSG |
| IV | 383 | 397 | PITKCQYSKPENCRL |

Using the 2VWD crystal structure from the Protein Data Base (PDB) we determined that regions 3 and 4 above are peptides that are highly surface exposed (Fig.2); while we show the peptides on one monomer of the dimeric structure, it is clear that those are not covered by the neighboring protein. Unfortunately, since the PDB structure starts from aa number 187, we could not visually demonstrate that regions 1 and 2 are also surface exposed, but on the basis of the ASA profile graph of Fig.1 and the evidence with regions 3 and 4, we assume that regions 1 and 2 are solvent accessible, although we have no information as of now whether the neighboring protein will cover them or not.

**Figure 2.**
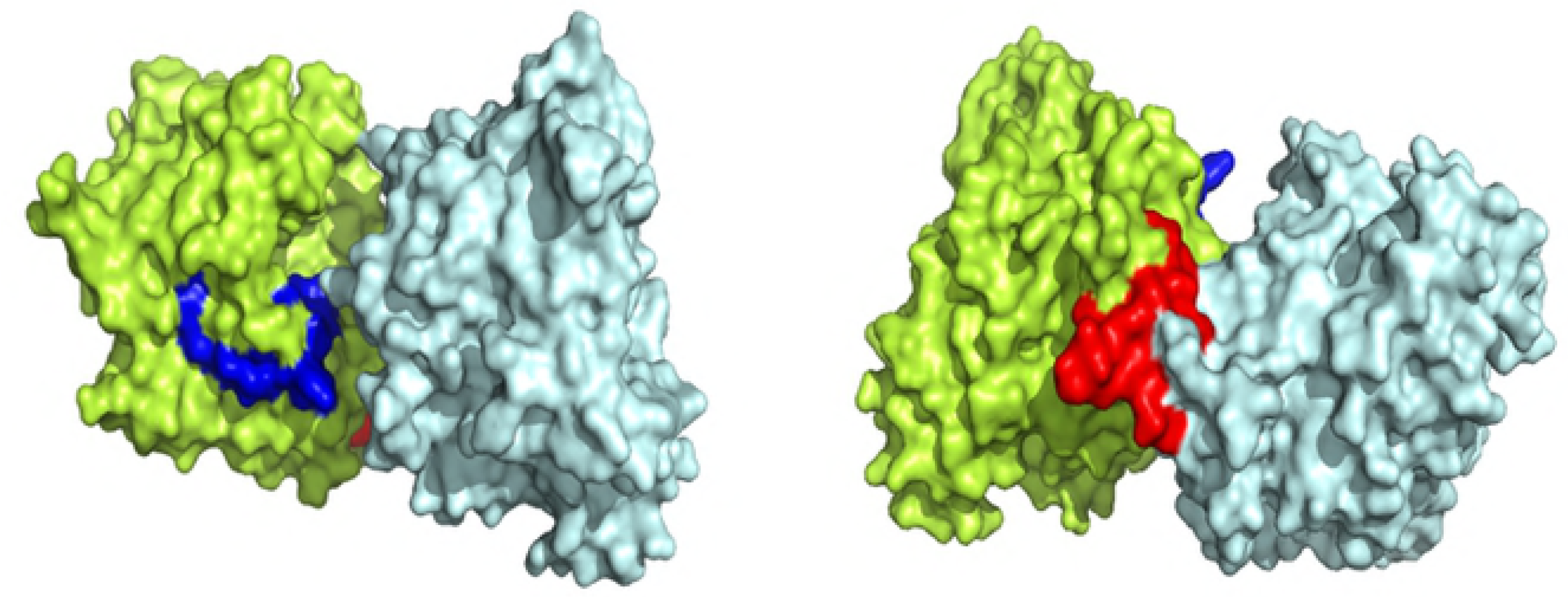
Display in space fill rendering of Nipah envelope protein structure (2VWD) by PyMol. The conserved surface exposed segments identified by comparison of protein variability and ASA profile analysis shown in Fig.l and given in Table 1 are highlighted in the limon green colored monomer. The segment marked in blue is the third region spanning aa numbers 301 to 315; the picture on the right is for region 4 spanning aa numbers 383-397 marked in red.

The next step was to search for existence of linear and conformational epitope regions within the identified segments so that we can in the current instance identify such epitopes and expect suitable adjuvants to enhance the immune response [17]. Our interest being generation of antibody response to the invading pathogens, we concentrate on MHC Class II molecules that mediate establishment of humoral immunity.

Using the IEDB server to determine the binding affinities for Human Leukocyte Antigens (HLA) for the four identified regions on the Nipah and Hendra virus glycoprotein sequence, the HLA alleles were chosen to provide a coverage of around 90% of the target population. Given the differences in disease severity in Malaysia and Bangladesh infections, we chose to use specific viral strains and host HLA alleles for each country, viz., Malaysia and Bangladesh, and we also computed the same for India (Kerala) population because of the recent Nipah virus outbreak in Kerala. Thus, we used the G glycoprotein from the AY029767 (Nipah virus isolate UMMC1, Malaysia), JN808863 (Nipah virus isolate NIVBGD2008RAJBARI, Bangladesh) and FJ513078 (Nipah virus isolate Ind-Nipah-07-FG, India) viral sequences for the three countries and their respective population HLA alleles.

All predictions were done using IEDB recommended procedure; most of the results reported are indicated to be consensus of the various measures used. The list of the binding affinities for MHC Class II T-cell epitopes, with percentile rank where low rank implies higher binding affinity, are given in Tables 2a and 2b; percentile rank of 10% and below are considered good binding strength. The default peptide length used by IEDB for binding strength computations is 15 residues. We select the peptides encompassing the identified group (Table 1) with a latitude of 4 to 5 residues to ensure good binding with the HLA alleles while remaining within the requirements of reasonably well conserved regions and high ASA; references to the crystal structure ensures the peptides are still surface exposed even if they are slightly adjusted from the regions as per Table 1.

**Table 2a.**
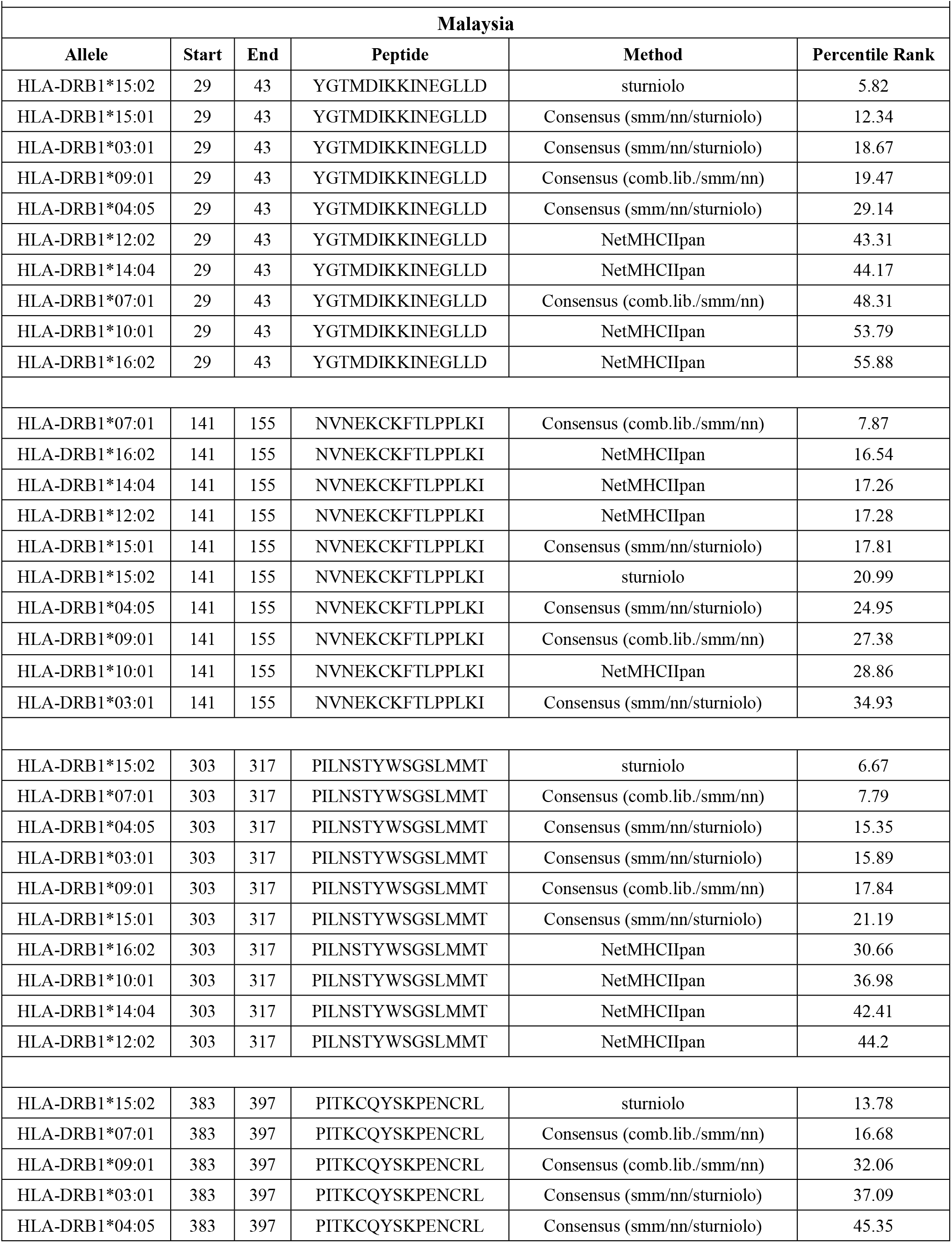
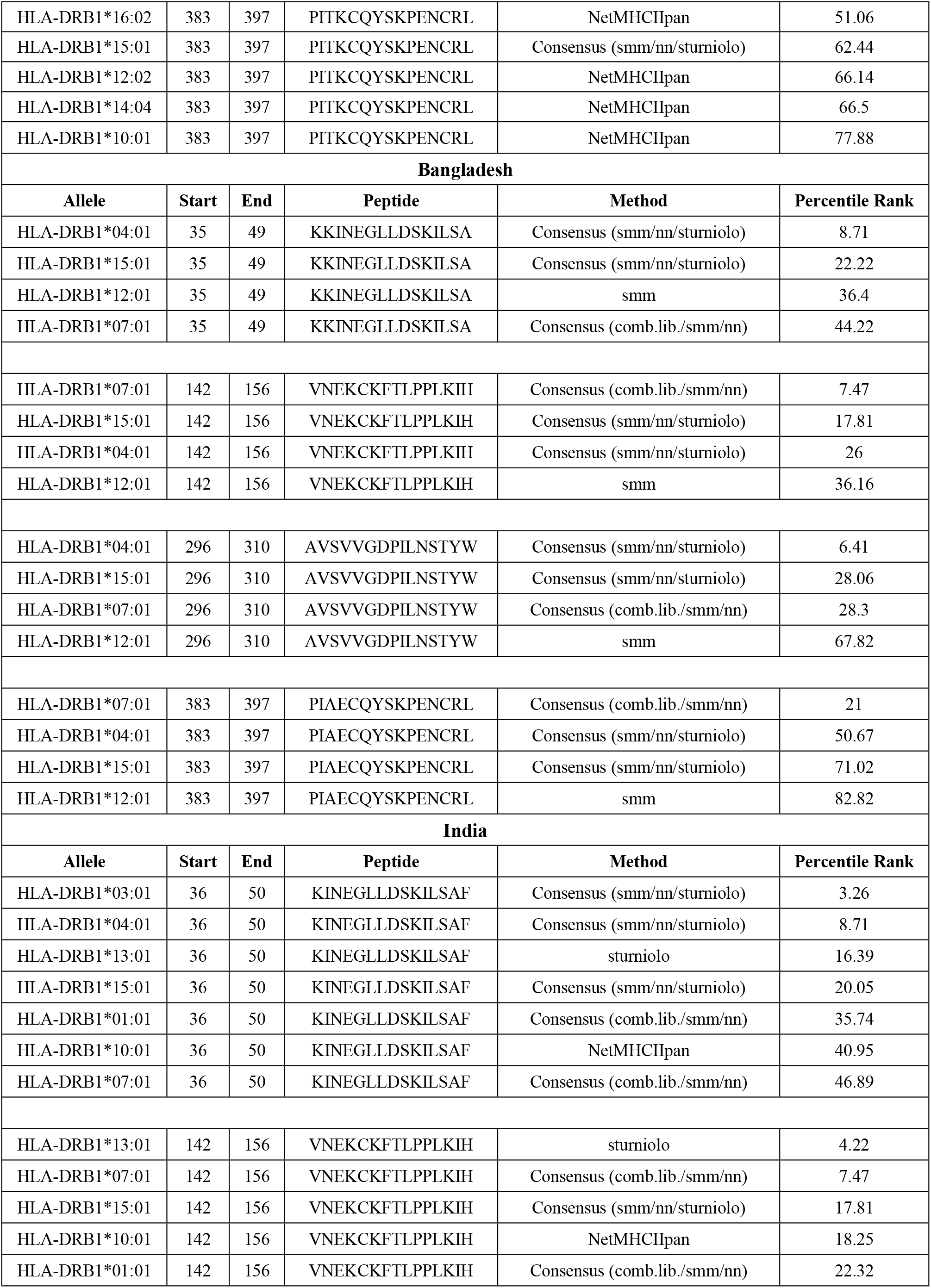
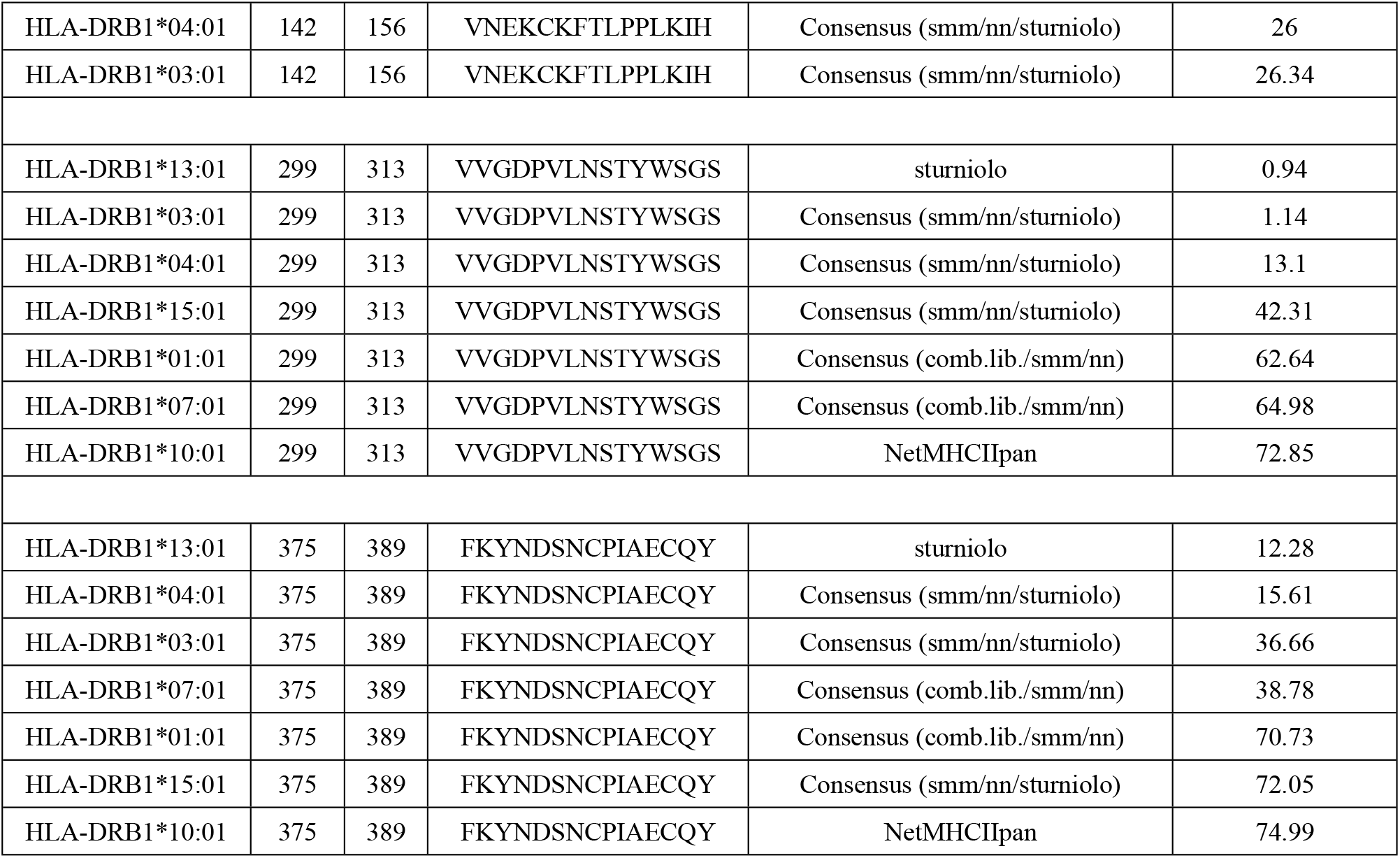
IEDB predictions of binding affinity for MHC-II HLA-DRB alleles of viral G-glycoprotein sequences from Malaysia, Bangladesh and India. The binding affinity is considered high for low percentile rank.

**Table 2b:**
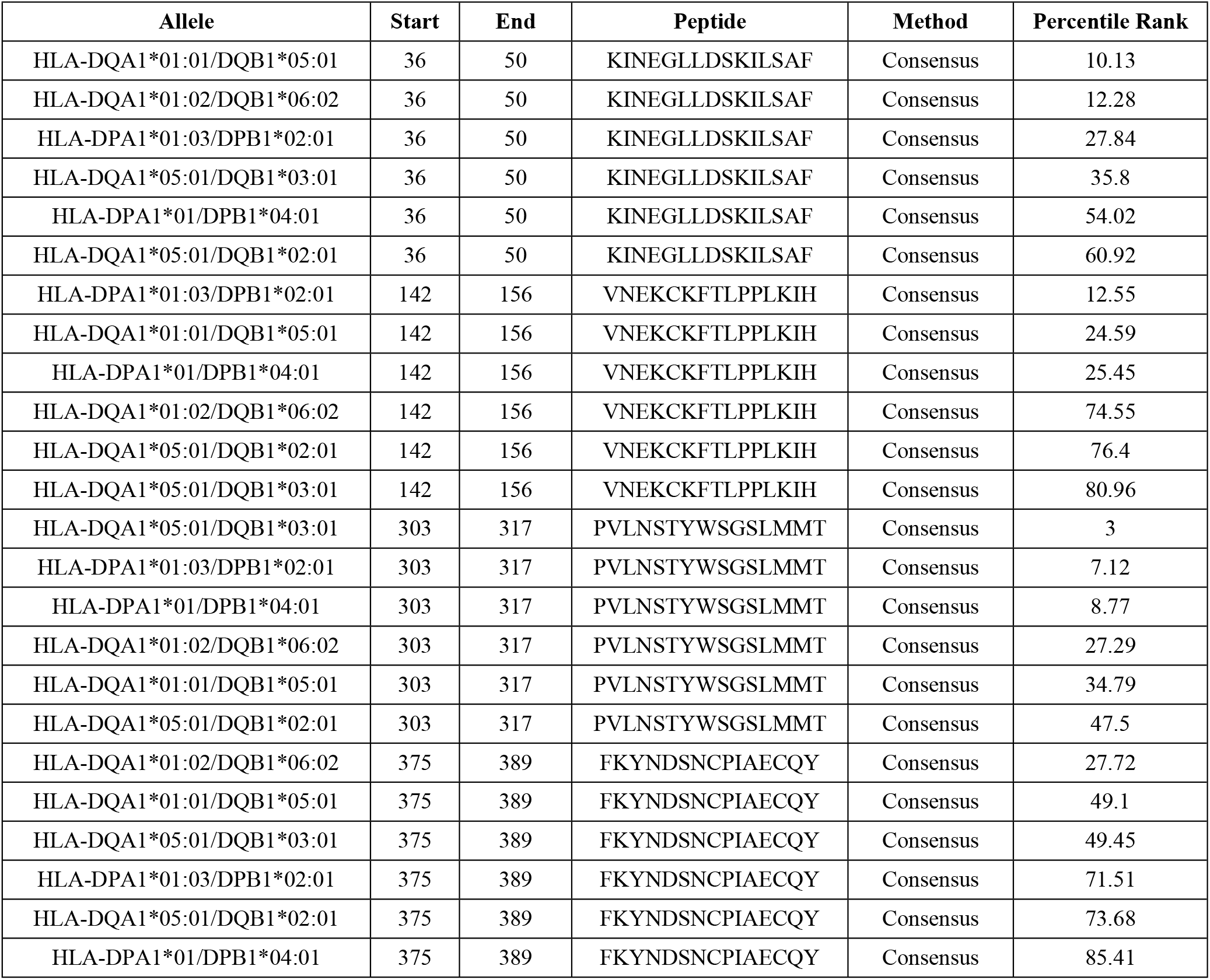
HLA DP/DQ alleles of the Indian population and their binding efficiencies with the Indian strain of the Nipah virus FJ513078 from IEDB.

The HLA-DRB alleles, which are the dominant types, are shown in Table 2a for the three countries/communities of Malaysia, Bangladesh and India. The Malaysian HLA alleles show comparatively large diversity to cover the population, Bangladesh the least while India falls in between. All regions show one or more alleles with good binding, region 4 just outside the 10-percentile rank for all 3 countries. All peptides of regions 3 and 4 (aa numbers 300+) were found to be surface exposed except for last two residues of Malaysian region 3 peptide (Table 2a). The peptides sequences are similar by construction, but are slightly divergent depending on the HLA profile and the prevailing viral types of the country. Thus, region 2 is almost the same for all 3 countries except for one residue displacement for Malaysia, but for region 2 we have a 6-residue displacement there compared to India and Bangladesh viruses. Similarly, region 3 has considerable overlaps, but in region 4 the India sample has only 7 residues in common with Bangladesh viral sequence and variation there also in case of the Malaysian sample.

The HLA DP/DQ allele profiles are available for India but not for Bangladesh and Malaysia. The results for Indian HLA alleles as determined through IEDB are given in Table 2b for the peptide segments seen in Table 2a except for a small displacement in region 3. This region showed the best results for the DP/DQ alleles amongst all the regions. The results for the other regions are the best available for the surface exposed conserved segments we are interested in.

As a further check on our selection of epitopes for the four regions of the viruses from the three countries, we determined the B-cell binding efficiencies using the ABCpred server. The summary results are given in Table 3 which shows that among the high binding efficiency epitopes predicted by ABCpred, all our determined epitope regions occur for all countries with high scores indicating good epitope potential.

**Table 3.**
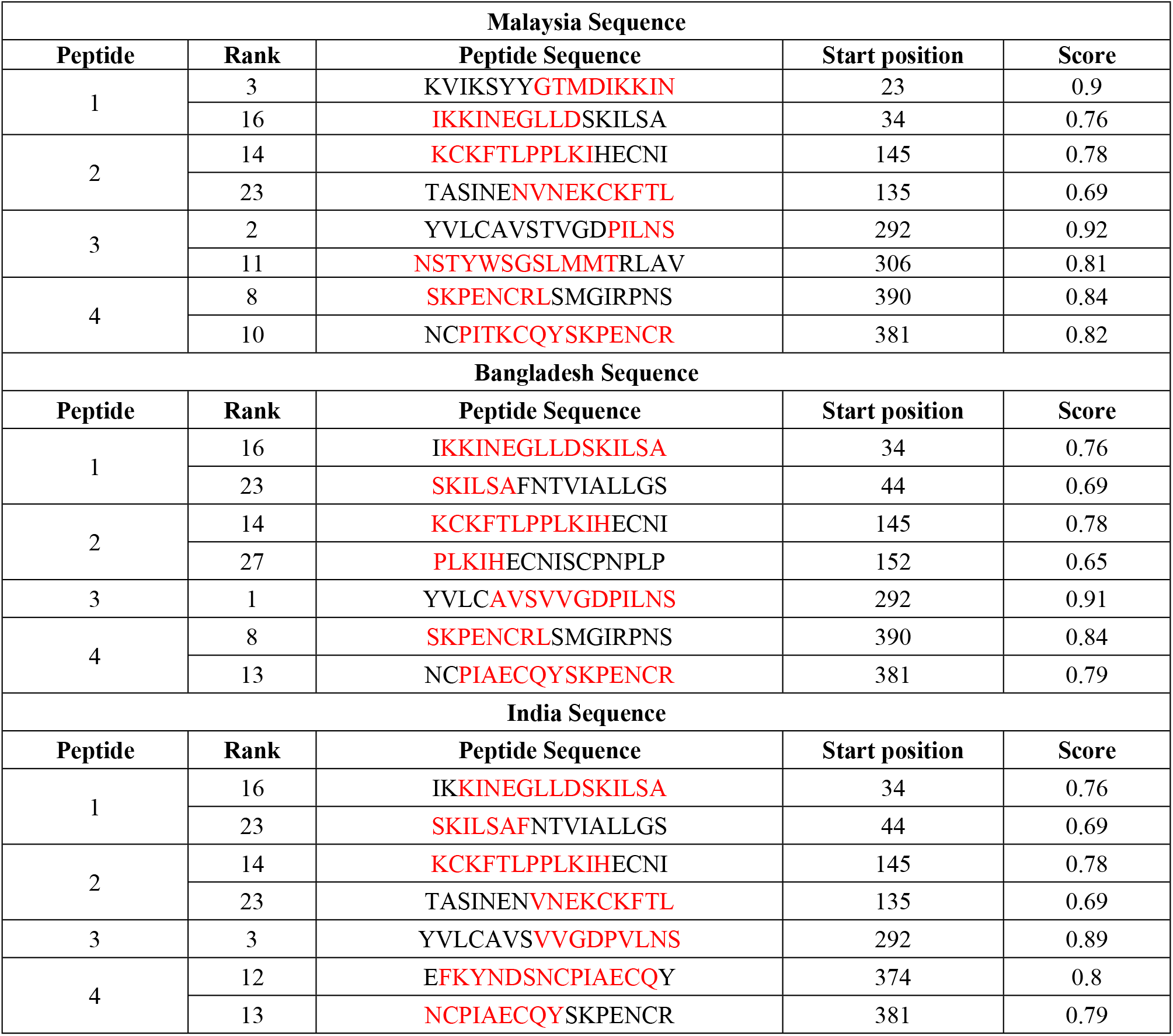
B-cell epitopes for selected peptides from ABCpred server. Low rank, high score indicates high binding affinity (cut-off score = 0.51). The red marks indicate the amino acids present in our desired peptide stretches.

The epitopes determined above are linear epitopes. It is known that viral epitopes are generally conformational; e.g., the West Nile virus epitopes are mostly non-linear and conformational as determined in mice models [37]. For Nipah and Hendra virus we also determine the binding affinities for conformational epitopes, through an Ellipro analysis of IEDB on the whole protein using the 2VWD Protein Data Bank data to determine presence of discontinuous epitopes for B-cell. The results are shown in Table 4; the parts of the sequences that match with our identified conserved segments are again marked in red. Thus, the peptide segments we had identified from the set of 35 sequences of the Nipah and Hendra glycoprotein appear to hold reasonable potential to respond as peptide vaccines.

**Table 4:**
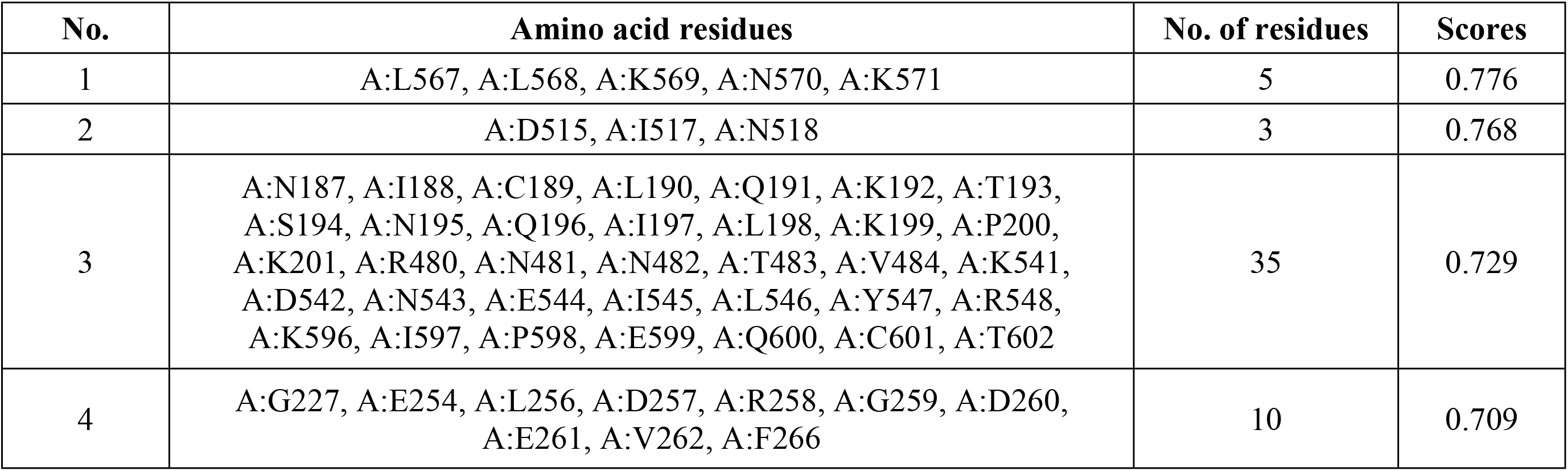
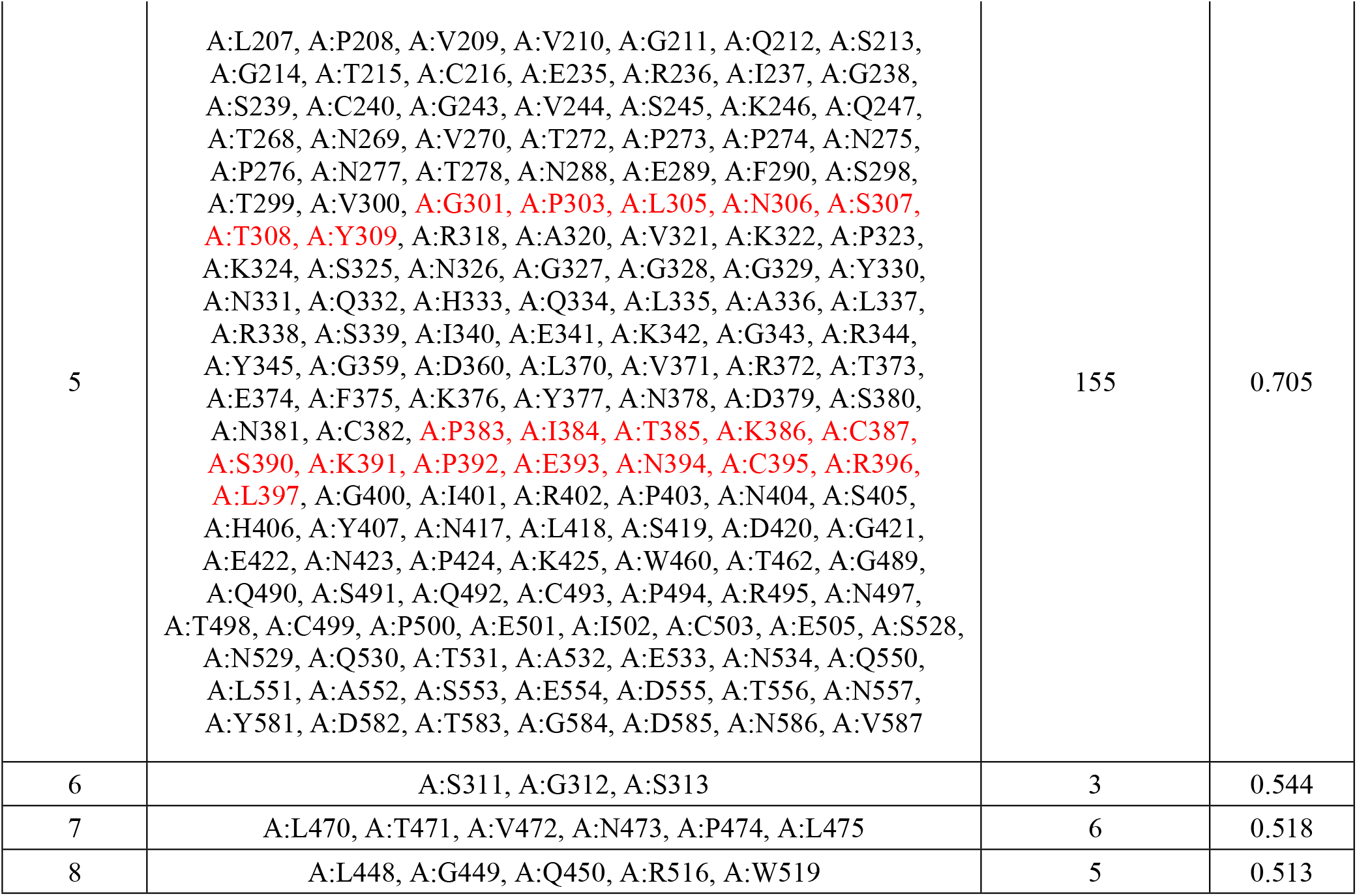
IEDB Ellipro predicted discontinuous epitope(s) for glycoprotein of Nipah and Hendra virus. The parts of the sequences that match with our identified conserved segments are marked in red.

| No. | Amino acid residues | No. of residues | Scores |
| --- | --- | --- | --- |
| 1 | A:L567, A:L568, A:K569, A:N570, A:K571 | 5 | 0.776 |
| 2 | A:D515, A:I517, A:N518 | 3 | 0.768 |
| 3 | A:N187, A:I188, A:C189, A:L190, A:Q191, A:K192, A:T193,<br>A:S194, A:N195, A:Q196, A:I197, A:L198, A:K199, A:P200,<br>A:K201, A:R480, A:N481, A:N482, A:T483, A:V484, A:K541,<br>A:D542, A:N543, A:E544, A:I545, A:L546, A:Y547, A:R548,<br>A:K596, A:I597, A:P598, A:E599, A:Q600, A:C601, A:T602 | 35 | 0.729 |
| 4 | A:G227, A:E254, A:L256, A:D257, A:R258, A:G259, A:D260,<br>A:E261, A:V262, A:F266 | 10 | 0.709 |
| 5 | A:L207, A:P208, A:V209, A:V210, A:G211, A:Q212, A:S213, A:G214, A:T215, A:C216, A:E235, A:R236, A:I237, A:G238, A:S239, A:C240, A:G243, A:V244, A:S245, A:K246, A:Q247, A:T268, A:N269, A:V270, A:T272, A:P273, A:P274, A:N275, A:P276, A:N277, A:T278, A:N288, A:E289, A:F290, A:S298, A:T299, A:V300, A:G301, A:P303, A:L305, A:N306, A:S307, A:T308, A:Y309, A:R318, A:A320, A:V321, A:K322, A:P323, A:K324, A:S325, A:N326, A:G327, A:G328, A:G329, A:Y330, A:N331, A:Q332, A:H333, A:Q334, A:L335, A:A336, A:L337, A:R338, A:S339, A:I340, A:E341, A:K342, A:G343, A:R344, A:Y345, A:G359, A:D360, A:L370, A:V371, A:R372, A:T373, A:E374, A:F375, A:K376, A:Y377, A:N378, A:D379, A:S380, A:N381, A:C382, A:P383, A:I384, A:T385, A:K386, A:C387, A:S390, A:K391, A:P392, A:E393, A:N394, A:C395, A:R396, A:L397, A:G400, A:I401, A:R402, A:P403, A:N404, A:S405, A:H406, A:Y407, A:N417, A:L418, A:S419, A:D420, A:G421, A:E422, A:N423, A:P424, A:K425, A:W460, A:T462, A:G489, A:Q490, A:S491, A:Q492, A:C493, A:P494, A:R495, A:N497, A:T498, A:C499, A:P500, A:E501, A:I502, A:C503, A:E505, A:S528, A:N529, A:Q530, A:T531, A:A532, A:E533, A:N534, A:Q550, A:L551, A:A552, A:S553, A:E554, A:D555, A:T556, A:N557, A:Y581, A:D582, A:T583, A:G584, A:D585, A:N586, A:V587 | 155 | 0.705 |
| 6 | A:S311, A:G312, A:S313 | 3 | 0.544 |
| 7 | A:L470, A:T471, A:V472, A:N473, A:P474, A:L475 | 6 | 0.518 |
| 8 | A:L448, A:G449, A:Q450, A:R516, A:W519 | 5 | 0.513 |

The peptide sequences we determined to be conserved surface exposed with good epitope potential were all tested for auto-immune threats by doing a protein-protein BLAST but no homology was found for any human proteins. Table 5 summarizes the 15-mer peptide results for these epitopes and could form a basis for eventual design of peptide vaccines, which ideally could be different for the different countries analyzed here.

**Table 5.**
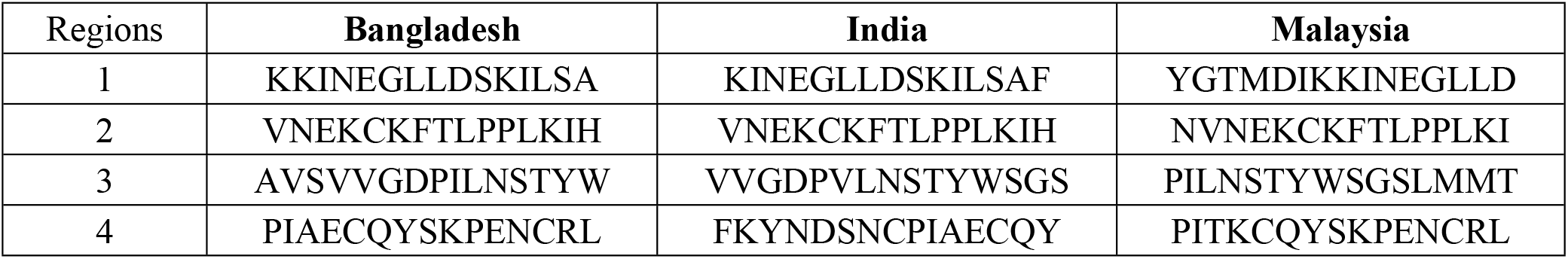
Summary of peptide vaccine epitopes.

## Summary and Conclusion

Thus, the results of our bioinformatics analyses reported above show that there are sufficient grounds for considering peptide vaccines as anti-viral agents against the Nipah and Hendra viruses. However, no peptide vaccines have been licensed as yet for human use, and the gap between the in silico designed vaccine candidates in the computer and the doctor’s use of the real vaccine in patients as the preventive tool against the viral disease is a long and tortuous one. In the case of the Zika virus where world attention was rivetted due to deformed births there have been many peptide vaccines in various phases of trials, but as yet no definitive conclusions have been arrived at. One of the difficulties in getting large scale trials going is that the epidemic has petered out [15]. Similarly, in the case of Nipah where the illnesses have been sporadic and localized, such trials would be even more difficult to conduct, no matter how important this step is for licensure. Trials with animal models remain the only recourse as a preparatory measure against a future need. While such trials have been reported for different types of vaccines against the Nipah virus as mentioned in the Introduction, these are all “one size fits all” type of vaccines. What we are advocating here are peptide vaccines that can be tailor-made to suit individual countries’ populations, which will be entirely possible on a “manufacturing” kind of scenario, something that is not possible, or would take an inordinately long time, with live-attenuated or VLP vaccines. If nothing else, peptide vaccines should be tested and got ready to combat any flare-up in Nipah virus infections given the human-to-human transmissibility of the virus and the extremely high CFRs that have been seen in India and Bangladesh so far.

## Acknowledgement

We acknowledge Prof Sukhen Das for his academic support during our research work.

